# Evolution in action: Brown-headed Cowbirds (*Molothrus ater*) adapt songs to 50 years of Ohio urbanization

**DOI:** 10.1101/2023.11.03.565548

**Authors:** Kristen Ruscitelli, Kaiya Provost

## Abstract

Technological advances in the last century have allowed humans to rapidly alter the landscape, resulting in a global loss of biodiversity. Of such alterations, is the conversion of rural land into urban space. In birds, urbanization has been found to impact song to facilitate improved auditory communication between conspecifics in areas with increased background noise. Given that birdsong is an important reproductive behavior, it is possible that urbanization promotes the evolution of birdsong through differences in fitness between individuals in urban acoustic environments. Here, we investigated if urbanization across Ohio impacts song characteristics of Brown-headed Cowbirds (*Molothrus ater*), a brood parasite associated with cropland and other altered habitats. To assess change in *M. ater* song over space and time, we compared syllable shape and syllable properties with metrics of urbanization. It was found that *M. ater* song differs significantly across geographic space and time, and is associated with changes in urbanization. Cowbird song of conspecifics in preferred habitat, such as cropland, demonstrates different syllable shape than conspecifics in urban space. This suggests that urbanization could facilitate change in *M. ater* song over time. These results provide additional insight into how anthropogenic habitat alteration facilitates cultural evolution in birds, and allows for increased understanding of how human behavior contributes to altered ecological interactions.

---

The Anthropocene is characterized by rapid environmental change caused by human activity, including the conversion of rural environment into an urban landscape (Lewis and Maslin 2015). Urbanization has a substantial impact on avian ecology through habitat loss and fragmentation (Crooks et al. 2004). Additionally, novel selective pressures associated with urban environments, such as human disturbances, artificial light pollution, and noise pollution, drive phenotypic change, have been demonstrated to lead to distinct differences in morphology and behavior between rural and urban conspecifics (Isaksson, 2018). For example, house finches (*Carpodacus mexicanus*) in both urban and non-urban habitats have been shown to experience strong, divergent, habitat-dependent selection on bill morphologies, despite close geographical proximity (Badyaevet al. 2008). This suggests that local adaptations can be maintained at small spatial scales in highly mobile species, such as songbirds. Additionally, differences in sexual selection between urban and non-urban habitats may lead to prezygotic isolation in dark-eyed juncos (*Junco hyemalis*), further suggesting an evolutionary distinction between urban and non-urban songbirds (Yeh, 2004). In a similar mechanism to that of genetic evolution, cultural evolution in birds facilitates the development of local song dialects that differ between populations (Aplin, 2019).

Given the substantial differences between urban and non-urban habitat, ornithologists have predicted that variation in birdsong should evolve between urban and non-urban conspecifics (Slabbekoorn and Boer-Visser, 2006). Variation in noise level and habitat structure between habitats results in differing levels of information transfer through birdsong (Dowlinget al. 2012). Characteristics of a habitat’s acoustic profile, such as attenuation, degradation, and acoustic masking are influenced by physical properties of that habitat (Brumm and Naguib, 2009). Given the distinct physical properties of urban and non-urban habitats, it is expected that each habitat has a distinct acoustic profile. Acoustic adaptation theory posits that characteristics of song that maximize transmission in a given habitat will be selected for, providing a basis for the claim that birdsong of urban and non-urban conspecifics will differ; consistent with this, birds in urban and non-urban habitats have observable differences in song that are thought to be driven by the increased background noise present in cities (Boncoraglio and Saino, 2007). For example, White-crowned Sparrows (*Zonotrichia leucophrys*) demonstrate faster trills and song with narrower bandwidth than rural conspecifics (Phillipset al. 2020). Additionally, common blackbirds (*Turdus merula*) have been shown to sing with higher-frequency elements at higher intensities in cities, potentially to reduce acoustic masking by urban noise (Nemethet al. 2013). Such differences have the potential to lead to reproductive isolation between urban and non-urban bird populations through a variety of mechanisms (Croninet al. 2022).

Territorial male birds respond differently to familiar versus unfamiliar song dialects (e.g., Lemon 1966). Males tend to react more strongly to familiar dialects than foreign dialects, potentially leading to males being less successful in establishing territory in areas with increased foreign dialects, and therefore reducing gene flow between populations (Slabbekoorn and Smith, 2002). Additionally, as with morphological traits, variation in species-specific song influences sexual selection in birds (Yeh, 2004). For example, females have been shown to prefer songs from males of local populations over songs from males from distant populations (Nowicki and Searcy, 2005). Given these factors, variation in dialect between populations may contribute to assortative mating. Brood parasites, specifically indigobirds (*Vidua chalybeata*), have demonstrated reproductive divergence resulting from song variation (Slabbekoorn and Smith, 2002). Seeing as local dialect preferences may lead to speciation, and birds in urban and non-urban environments have observable differences in song, there is the possibility that differences in dialect between urban and non-urban conspecifics contribute to reproductive divergence. Further examination of the relationship between urbanization and change in brood parasite song facilitates an increased understanding of the mechanism by which anthropogenic habitat alteration contributes to ecological change, such as speciation.

## Focal System

Brown-headed Cowbirds (*Molothrus ater*) are a species of brood parasite that are commonly found throughout the state of Ohio and their habitat typically consists of open land, such as cropland or grazing land (Lowther 2020). Over the last half-century, open habitat has been increasingly converted into urban landscape, providing the opportunity to examine the association between such changes in habitat and changes in *M. ater* song over approximately 50 years (Elliset al. 2020). Given previous literature findings that suggest differences in birdsong between urban and non-urban conspecifics, it is expected that there has been an observable change in *M. ater* song over the last half-century that is associated with a decrease in open habitat and an increase in urbanization. If such prediction is indeed observed, it would provide additional insight into how anthropogenic habitat alteration may contribute to change in ecological dynamics in brood parasites.

Here, we examined change in *M. ater* song properties and shape in relation to anthropogenic habitat alteration over approximately the last 50 years. We predict that these changes in *M. ater* song would be correlated with increased urbanization to accommodate the shifting acoustic landscape in urban settings. To test this prediction, recordings of *M. ater* song were annotated and song summary statistics were collected. Environmental data was obtained from the Anthromes dataset to determine the degree of change in urbanization across Ohio over the last half-century. Statistical analyses were then conducted to determine if change in *M. ater* song significantly differs across time, geographic location, and degree of urbanization.

## Methods

### Sound Data

*Molothrus ater* song recordings (*n*=86) were accessed from the Borror Laboratory of Bioacoustics database at Ohio State University (Table 1). These recordings were made over the course of some ∼52 years (1948-2000) (see supplemental). Song data was exported to Raven Pro (version 1.6.4) and recordings were annotated such that individual syllables were separated into individual selections. Raven Pro was used to calculate a suite of bioacoustic summary statistics (see Table 2) that described frequency, time, duration, bandwidth, slope, and other aspects of the individual syllables. With these summary statistics we performed a principal components analysis (PCA) to reduce the dimensionality of the data, particularly because many of the summary statistics were highly correlated. We performed the PCA using the prcomp function in the stats package in R version 4.3.1 (R Core Team 2023) on 55 summary statistics that were numeric, had no missing data, and were not all the same value. These data were centered and scaled before the PCA was performed. The resulting principal components (PCs) are referred to as “Prop.PCs” henceforth. We analyze the first five PCs of this analysis (see Results).

**Table 1:**
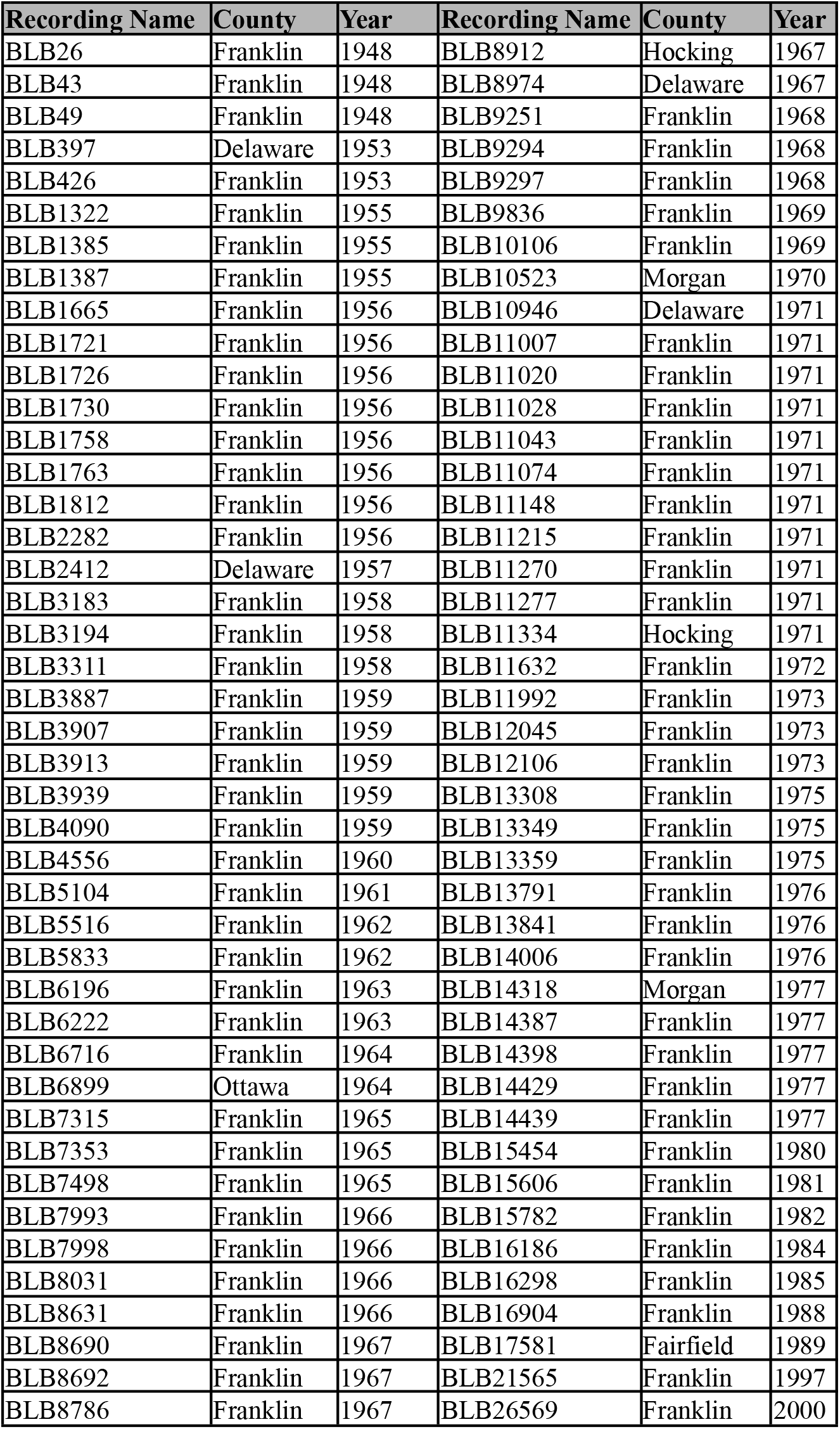
*Molothrus ater* recordings used across Ohio. Note that recordings are split across two halves of the table.

**Table 2:** principal components analysis on 55 bioacoustic summary statistics. Relative importance of each PC is shown.

| Principal component | Standard deviation | Proportion of variance | Cumulative proportion |
| --- | --- | --- | --- |
| Prop.PC1 | 4.243 | 0.327 | 0.327 |
| Prop.PC2 | 3.597 | 0.235 | 0.562 |
| Prop.PC3 | 2.452 | 0.109 | 0.672 |
| Prop.PC4 | 1.988 | 0.072 | 0.744 |
| Prop.PC5 | 1.847 | 0.062 | 0.806 |
| Prop.PC6 | 1.603 | 0.047 | 0.852 |
| Prop.PC7 | 1.173 | 0.025 | 0.877 |
| Prop.PC8 | 1.064 | 0.021 | 0.898 |
| Prop.PC9 | 1.001 | 0.018 | 0.916 |
| Prop.PC10 | 0.861 | 0.013 | 0.930 |
| Prop.PC11 | 0.804 | 0.012 | 0.941 |
| Prop.PC12 | 0.738 | 0.010 | 0.951 |
| Prop.PC13 | 0.678 | 0.008 | 0.960 |
| Prop.PC14 | 0.632 | 0.007 | 0.967 |
| Prop.PC15 | 0.523 | 0.005 | 0.972 |
| Prop.PC16 | 0.480 | 0.004 | 0.976 |
| Prop.PC17 | 0.442 | 0.004 | 0.980 |
| Prop.PC18 | 0.435 | 0.003 | 0.983 |
| Prop.PC19 | 0.383 | 0.003 | 0.986 |
| Prop.PC20 | 0.351 | 0.002 | 0.988 |
| Prop.PC21 | 0.343 | 0.002 | 0.990 |
| Prop.PC22 | 0.326 | 0.002 | 0.992 |
| Prop.PC23 | 0.313 | 0.002 | 0.994 |
| Prop.PC24 | 0.297 | 0.002 | 0.995 |
| Prop.PC25 | 0.256 | 0.001 | 0.997 |
| Prop.PC26 | 0.235 | 0.001 | 0.998 |
| Prop.PC27 | 0.197 | 0.001 | 0.998 |
| Prop.PC28 | 0.158 | 4.500E-04 | 0.999 |
| Prop.PC29 | 0.146 | 3.900E-04 | 0.999 |
| Prop.PC30 | 0.135 | 3.300E-04 | 0.999 |
| Prop.PC31 | 0.132 | 3.200E-04 | 1.000 |
| Prop.PC32 | 0.094 | 1.600E-04 | 1.000 |
| Prop.PC33 | 0.052 | 5.000E-05 | 1.000 |
| Prop.PC34 | 0.005 | 0 | 1.000 |
| Prop.PC35 | 0.003 | 0 | 1.000 |
| Prop.PC36 | 0.002 | 0 | 1.000 |
| Prop.PC37 | 0.001 | 0 | 1.000 |
| Prop.PC38 | 0.001 | 0 | 1.000 |
| Prop.PC39 | 4.110E-05 | 0 | 1.000 |
| Prop.PC40 | 3.980E-06 | 0 | 1.000 |
| Prop.PC41 | 1.280E-06 | 0 | 1.000 |
| Prop.PC42 | 1.010E-06 | 0 | 1.000 |
| Prop.PC43 | 9.720E-07 | 0 | 1.000 |
| Prop.PC44 | 8.030E-07 | 0 | 1.000 |
| Prop.PC45 | 3.310E-07 | 0 | 1.000 |
| Prop.PC46 | 2.180E-07 | 0 | 1.000 |
| Prop.PC47 | 7.900E-08 | 0 | 1.000 |
| Prop.PC48 | 7.320E-08 | 0 | 1.000 |
| Prop.PC49 | 1.880E-15 | 0 | 1.000 |
| Prop.PC50 | 7.380E-16 | 0 | 1.000 |
| Prop.PC51 | 7.330E-16 | 0 | 1.000 |
| Prop.PC52 | 3.850E-16 | 0 | 1.000 |
| Prop.PC53 | 3.850E-16 | 0 | 1.000 |
| Prop.PC54 | 3.850E-16 | 0 | 1.000 |
| Prop.PC55 | 2.870E-16 | 0 | 1.000 |

We used the SoundShape package (Rocha and Romano 2021) to extract shape data from our selections as well. This package normalizes syllables to make them directly comparable with respect to shape using geometric morphometric principles. Our syllables had 14,400 points each. We then performed a PCA on the resulting shape data, whose resulting PCs are referred to as “Shp.PCs” henceforth. We analyze the first five PCs of this analysis (see Results).

### Environmental Data

To extract data about the urbanization of Ohio across the decades, we used the Anthromes dataset (Elliset al. 2020) for the years 1940 and 2017. Anthromes are categorizations of the urbanization levels from underlying data about the percent cropland, percent land used for grazing, percent land used to irrigate rice fields, total percent of land that is irrigated, the census population, and the percent of urban occupancy. Rather than using the categorical data in the Anthromes dataset, we directly analyzed the raw data from each category because cropland and grazing land are assumed to be critical for cowbirds.

For each recording, we extracted the corresponding Anthromes values at the latitude and longitude of the recording for the years 1940 and 2017. We then calculated the difference in values between those two years as a measure of the change in urbanization using the raster version 3.6-14 package in R (Hijmans 2023). We therefore had a measure of the amount of change in cropland, grazing land, and the other four measures of urbanization across the time period of the recordings.

### Analyses

To evaluate our hypotheses, we performed statistical tests using ANOVAs for categorical data and linear models for quantitative data. ANOVAs were performed using the aov function and linear models with the lm function, both in the stats package in R. For both linear models and ANOVAs, we used a significance cutoff value of alpha = 0.05. ANOVAs were conducted to compare the Prop.PCs and the Shp.PCs against variables such as County and Month Collected. A Tukey’s Honest Significant Differences test was then conducted to evaluate significant ANOVA results. These were performed using the TukeyHSD function in the stats package in R. Linear models were used to compare the Prop.PCs and Shp.PCs against variables such as Year Collected, the six urbanization metrics, Latitude, and Longitude. For some simpler comparisons, we also performed t-tests using the t.test function in the stats package in R. Our code is available at https://github.com/kaiyaprovost/bioacoustics/blob/main/cowbird_processing.R.

## Results

The 86 *M. ater* recordings consisted of 1,040 total syllables. These recordings came from four counties, three in central Ohio (Franklin, Delaware, Hocking counties) and one in northern Ohio on Lake Erie (Ottawa county) (Fig. 1). When we perform a PCA on the 55 summary statistics we extracted from our syllables, we find that the first two PCs explained 56.2% of the data and the first five explained 80.5% of the data (Table 2). For the summary statistics PCA, Prop.PC1 primarily describes when in the recording the syllable is located with respect to time, such that high Prop.PC1 means a syllable late in the recording. Syllables with high Prop.PC2 are high frequency and have lots of energy, entropy, and power. They also have long durations. Syllables with high Prop.PC3 have short durations and few inflection points, but are also high in frequency. Syllables with High Prop.PC4 have low bandwidth, low entropy, and low frequency, but high power and high slope. Syllables with high Prop.PC5 are short in duration and low frequency, but have high bandwidth (Table 3). When we perform a principal component analysis on the shape of our syllables, we find that the first 24 PCs explain 50.1% of the data, with the first three PCs explaining 18.1%, 5.6%, and 3.1%, respectively (Table 4).

**Figure 1:**
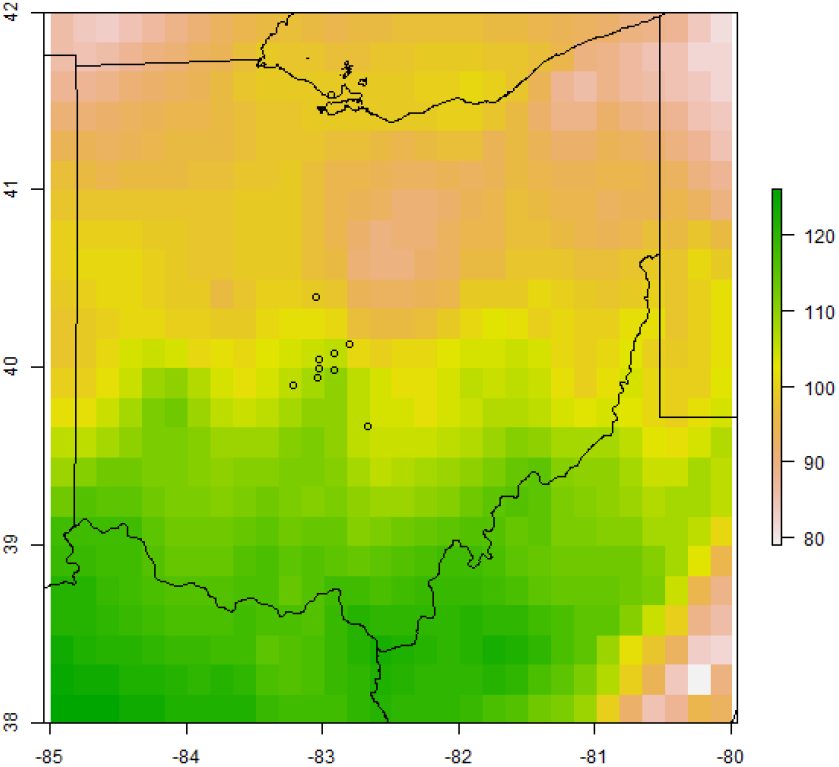
Cowbird recording locations across Ohio. Circles indicate individual recordings; note that multiple recordings are taken at the same locality. Background colors indicate mean annual temperature across Ohio.

**Table 3:** loading/rotation of bioacoustic summary statistics on the first five PCs, which explain ∼80% of the variation. PFC=peak frequency contour. Variables are sorted by PC1 value.

| Variable | PC1 | PC2 | PC3 | PC4 | PC5 |
| --- | --- | --- | --- | --- | --- |
| Average Power Density | -0.041 | 0.176 | 0.048 | 0.227 | 0.005 |
| Inband Power | -0.034 | 0.213 | 0.020 | 0.153 | 0.014 |
| Energy | -0.034 | 0.233 | -0.042 | 0.156 | 0.012 |
| Peak Power Density | -0.030 | 0.217 | -0.008 | 0.181 | 0.016 |
| Length of Frames | -0.023 | 0.166 | -0.277 | 0.146 | 0.024 |
| Sample Length | -0.023 | 0.166 | -0.277 | 0.146 | 0.024 |
| Duration 100% | -0.023 | 0.166 | -0.277 | 0.146 | 0.024 |
| Duration 90% | -0.022 | 0.144 | -0.297 | 0.135 | 0.000 |
| PFC Inflections | -0.021 | 0.143 | -0.300 | 0.131 | 0.025 |
| Duration 50% | -0.021 | 0.132 | -0.284 | 0.124 | -0.006 |
| Minimum Entropy | -0.020 | -0.079 | -0.081 | -0.251 | -0.028 |
| Maximum Entropy | -0.016 | 0.220 | -0.055 | -0.135 | 0.020 |
| Average Entropy | -0.015 | 0.033 | -0.220 | -0.268 | 0.045 |
| Aggregate Entropy | -0.014 | 0.066 | -0.163 | -0.367 | 0.021 |
| Center Relative Time | -0.012 | -0.026 | -0.114 | -0.049 | -0.471 |
| Time 95% Relative | -0.009 | -0.059 | -0.165 | -0.065 | -0.253 |
| Time 25% Relative | -0.009 | -0.025 | -0.042 | -0.025 | -0.423 |
| Time 5% Relative | -0.008 | 0.031 | 0.046 | -0.022 | -0.265 |
| Time 75% Relative | -0.007 | -0.037 | -0.152 | -0.047 | -0.403 |
| Peak Frequency Relative Time | -0.006 | -0.016 | -0.116 | -0.032 | -0.430 |
| Bandwidth 90% | -0.004 | 0.052 | -0.089 | -0.389 | 0.106 |
| PFC Maximum Slope | -0.002 | 0.150 | -0.112 | -0.195 | 0.098 |
| PFC Minimum Slope | 0.000 | -0.158 | 0.108 | 0.163 | -0.112 |
| Bandwidth 100% | 0.000 | 0.218 | -0.068 | -0.050 | 0.039 |
| Boundary Maximum Frequency | 0.001 | 0.240 | 0.003 | -0.027 | 0.004 |
| Bandwidth 50% | 0.002 | 0.020 | -0.073 | -0.336 | 0.074 |
| Boundary Minimum Frequency | 0.002 | 0.124 | 0.249 | 0.073 | -0.120 |
| Maximum Frequency | 0.002 | 0.230 | 0.143 | -0.076 | -0.080 |
| Peak Frequency | 0.002 | 0.230 | 0.143 | -0.076 | -0.080 |
| Frequency 90% | 0.004 | 0.220 | 0.099 | -0.222 | -0.003 |
| PFC Maximum Frequency | 0.005 | 0.242 | 0.061 | -0.115 | 0.022 |
| Frequency 25% | 0.007 | 0.231 | 0.171 | -0.010 | -0.089 |
| Frequency 75% | 0.007 | 0.232 | 0.135 | -0.151 | -0.054 |
| Frequency 5% | 0.008 | 0.219 | 0.192 | 0.058 | -0.092 |
| Center Frequency | 0.008 | 0.235 | 0.154 | -0.078 | -0.075 |
| PFC Minimum Frequency | 0.012 | 0.131 | 0.266 | 0.047 | -0.115 |
| PFC Average Slope | 0.016 | 0.019 | 0.009 | -0.079 | -0.074 |
| Selection Number | 0.221 | 0.010 | 0.013 | -0.023 | 0.009 |
| End Samples | 0.235 | 0.011 | -0.018 | 0.009 | -0.002 |
| End Time | 0.235 | 0.011 | -0.018 | 0.009 | -0.002 |
| End File Samples | 0.235 | 0.011 | -0.018 | 0.009 | -0.002 |
| End Clock Time | 0.235 | 0.011 | -0.018 | 0.009 | -0.002 |
| Time 95% | 0.235 | 0.010 | -0.017 | 0.009 | -0.003 |
| Time 75% | 0.235 | 0.010 | -0.017 | 0.009 | -0.003 |
| Maximum Time | 0.235 | 0.009 | -0.016 | 0.008 | -0.004 |
| Peak Frequency Time | 0.235 | 0.009 | -0.016 | 0.008 | -0.004 |
| Center Time | 0.235 | 0.009 | -0.016 | 0.008 | -0.003 |
| Time 25% | 0.235 | 0.009 | -0.015 | 0.008 | -0.003 |
| Time 5% | 0.235 | 0.009 | -0.014 | 0.007 | -0.003 |
| Begin Samples | 0.235 | 0.008 | -0.014 | 0.007 | -0.002 |
| Begin Time | 0.235 | 0.008 | -0.014 | 0.007 | -0.002 |
| Begin File Samples | 0.235 | 0.008 | -0.014 | 0.007 | -0.002 |
| Begin Clock Time | 0.235 | 0.008 | -0.014 | 0.007 | -0.002 |
| File Offset Time | 0.235 | 0.008 | -0.014 | 0.007 | -0.002 |
| Begin Date-Time | 0.235 | 0.008 | -0.014 | 0.007 | -0.002 |

**Table 4:** principal components analysis on shape metrics of individual syllables (n=14,400 individual points). Importance of the first 24 PCs are shown which explain ∼50% of the data.

| Principal component | Standard deviation | Proportion of variance | Cumulative proportion |
| --- | --- | --- | --- |
| Shp.PC1 | 62.679 | 0.182 | 0.182 |
| Shp.PC2 | 34.966 | 0.057 | 0.238 |
| Shp.PC3 | 25.785 | 0.031 | 0.269 |
| Shp.PC4 | 22.651 | 0.024 | 0.292 |
| Shp.PC5 | 21.570 | 0.022 | 0.314 |
| Shp.PC6 | 20.051 | 0.019 | 0.333 |
| Shp.PC7 | 19.293 | 0.017 | 0.350 |
| Shp.PC8 | 18.673 | 0.016 | 0.366 |
| Shp.PC9 | 17.175 | 0.014 | 0.379 |
| Shp.PC10 | 16.144 | 0.012 | 0.392 |
| Shp.PC11 | 15.659 | 0.011 | 0.403 |
| Shp.PC12 | 14.826 | 0.010 | 0.413 |
| Shp.PC13 | 14.559 | 0.010 | 0.423 |
| Shp.PC14 | 14.307 | 0.009 | 0.432 |
| Shp.PC15 | 13.639 | 0.009 | 0.441 |
| Shp.PC16 | 13.328 | 0.008 | 0.449 |
| Shp.PC17 | 12.766 | 0.008 | 0.457 |
| Shp.PC18 | 12.453 | 0.007 | 0.464 |
| Shp.PC19 | 12.241 | 0.007 | 0.471 |
| Shp.PC20 | 11.876 | 0.007 | 0.477 |
| Shp.PC21 | 11.788 | 0.006 | 0.484 |
| Shp.PC22 | 11.438 | 0.006 | 0.490 |
| Shp.PC23 | 11.294 | 0.006 | 0.496 |
| Shp.PC24 | 11.134 | 0.006 | 0.501 |

### Statistical analysis

*Molothrus ater* song differs significantly across the different counties (Table 5). For Prop.PC1 (recording length), Prop.PC4 (bandwidth and power), and all five Shp.PCs (Fig. 2A), syllables are significantly different, though the exact differences vary. Some of this differentiation is likely due to the fact that one county (Ottawa) is much further away from the others, being on Lake Erie. Other differences also include the fact that Franklin County is by far the most urbanized of the four, containing the capital region. Indeed, almost all of the differences detected are either due to Ottawa or Franklin being different.

**Figure 2:**
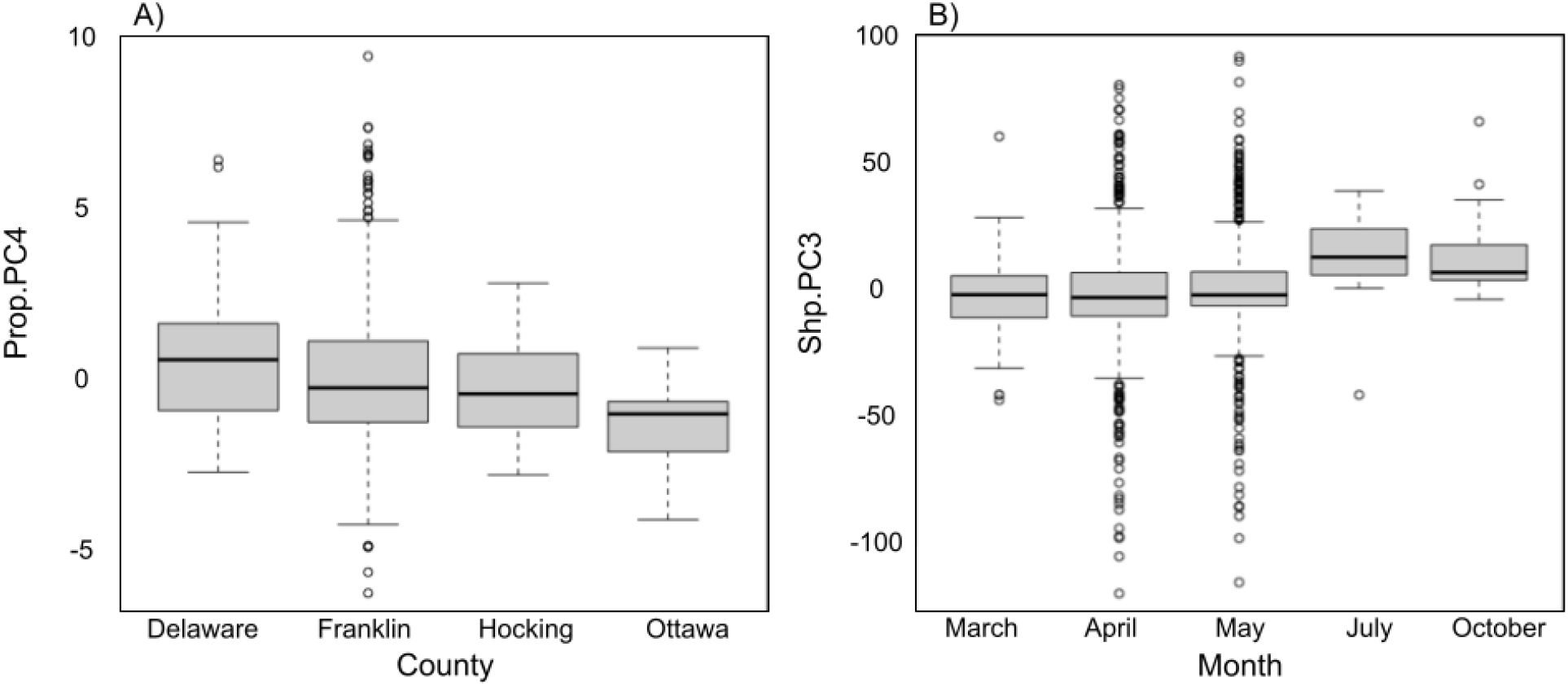
cowbird syllables vary according to county and month. a) song properties (Prop.PC4) significantly vary by county. b) song shapes (Shp.PC3) significantly vary by month.

**Table 5:** significant ANOVA and Tukey’s Honest Significant Difference results between predictor variables (county, month, year) and response variables (Shp.PCs and Prop.PCs). “Tukey” columns show p-values for the comparison indicated.

| Predictor | Response | DF | SSQ | MSQ | F | P | Tukey |  |  |
| --- | --- | --- | --- | --- | --- | --- | --- | --- | --- |
| Prop.PC1 | County | 3 | 604 | 201.38 | 11.54 | < 0.001 | Franklin-Delaware: < 0.001 | Ottawa-Franklin: 0.010 |  |
| Prop.PC4 | County | 3 | 39 | 12.845 | 3.27 | 0.020 | Ottawa-Delaware: 0.038 |  |  |
| Shp.PC1 | County | 3 | 161790 | 53930 | 14.25 | < 0.001 | Franklin-Delaware: < 0.001 | Ottawa-Delaware: 0.010 | Hocking-Franklin: 0.003 |
| Shp.PC2 | County | 3 | 21619 | 7206 | 5.98 | < 0.001 | Hocking-Delaware: < 0.001 | Ottawa-Hocking: 0.038 | Hocking-Franklin: 0.003 |
| Shp.PC3 | County | 3 | 7252 | 2417.2 | 3.66 | 0.012 | Ottawa-Hocking: 0.019 | Hocking-Franklin: 0.034 |  |
| Shp.PC4 | County | 3 | 15756 | 5252 | 10.52 | < 0.001 | Franklin-Delaware: < 0.001 | Hocking-Delaware: < 0.001 |  |
| Shp.PC5 | County | 3 | 10455 | 3485 | 7.63 | < 0.001 | Franklin-Delaware: < 0.001 | Hocking-Delaware: < 0.001 |  |
| Prop.PC3 | Month | 1 | 58 | 58.19 | 9.77 | 0.001 |  |  |  |
| Prop.PC4 | Month | 1 | 57 | 57.36 | 14.72 | < 0.001 |  |  |  |
| Shp.PC3 | Month | 1 | 3255 | 3255 | 6.38 | 0.011 |  |  |  |
| Shp.PC4 | Month | 1 | 9169 | 9169 | 13.96 | < 0.001 |  |  |  |
| Shp.PC5 | Month | 1 | 10708 | 10708 | 23.52 | < 0.001 |  |  |  |

Songs are also different across the months (Fig. 2B). Prop.PC3 (duration, frequency) and Prop.PC4 (bandwidth and power) show variation associated with month, as do Shp.PC3, Shp.PC4, and Shp.PC5. Differences across the months may be related to differences in breeding effort.

Change in cropland has a large impact on song both for song properties (Prop.PC1, recording length) and syllable shape (Shp.PC1, Shp.PC2, Shp.PC4, and Shp.PC5; Table 6; Fig. 3). This indicates that syllable shapes are different in regions that have dramatically lost or gained cropland compared to areas where they have not. Other aspects of the environment surveyed do not correlate at all with cowbird song.

**Figure 3:**
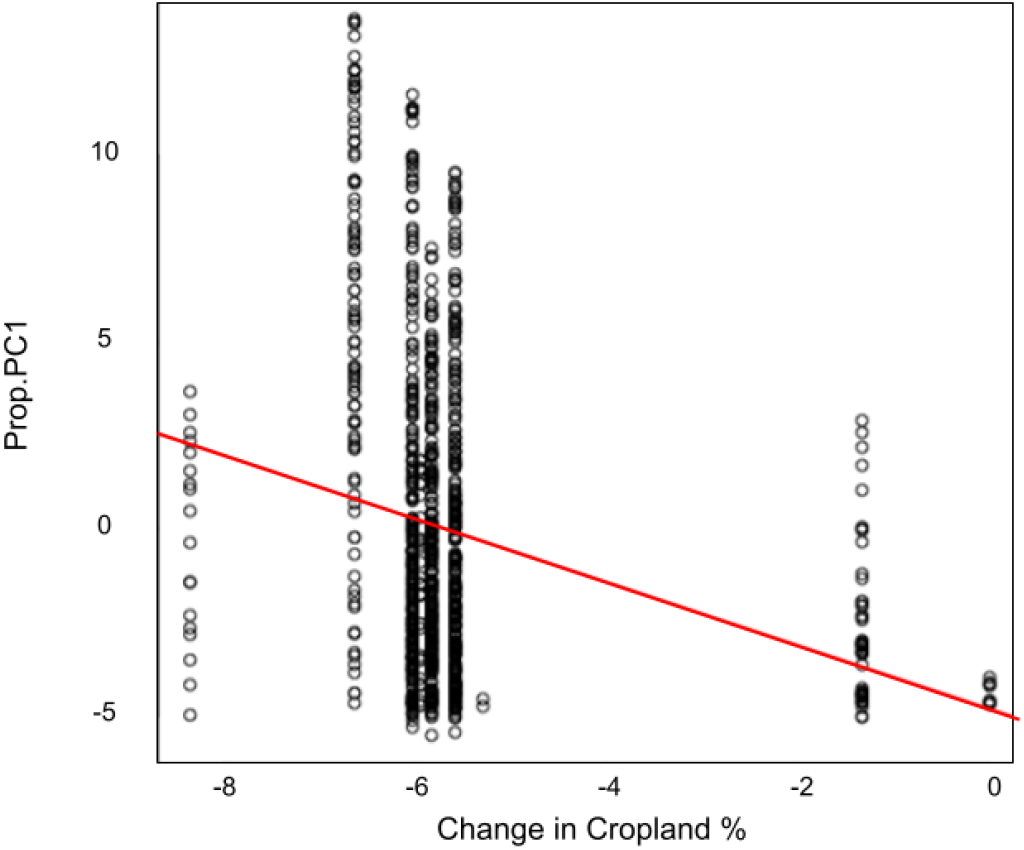
cowbird syllable properties (Prop.PC1) vary according to change in cropland percentage over time. Red line indicates a statistically significant relationship.

**Table 6:** significant linear model results between predictor variables (change in cropland, Latitude, Longitude, year) and response variables (Shp.PCs and Prop.PCs).

| Comparison | R<sup>2</sup> | Intercept | Int. Std. Err. | Int. T | Int. P | Coefficient | Coef. Std. Err. | Coef. T | Coef. P |
| --- | --- | --- | --- | --- | --- | --- | --- | --- | --- |
| cropland, Prop.PC1 | 0.056 | -4.805 | 0.629 | -7.635 | < 0.001 | -0.847 | 0.109 | -7.803 | < 0.001 |
| cropland, Shp.PC1 | 0.021 | -44.547 | 9.470 | -4.704 | < 0.001 | -7.849 | 1.634 | -4.804 | < 0.001 |
| cropland, Shp.PC2 | 0.005 | -12.926 | 5.326 | -2.427 | 0.015 | -2.277 | 0.919 | -2.479 | 0.013 |
| cropland, Shp.PC4 | 0.008 | 10.473 | 3.444 | 3.041 | 0.002 | 1.845 | 0.594 | 3.105 | 0.002 |
| cropland, Shp.PC5 | 0.010 | -10.904 | 3.277 | -3.328 | < 0.001 | -1.921 | 0.565 | -3.398 | < 0.001 |
| Latitude, Prop.PC1 | 0.023 | 110.917 | 22.209 | 4.994 | < 0.001 | -2.776 | 0.556 | -4.994 | < 0.001 |
| Latitude, Prop.PC4 | 0.015 | 42.694 | 10.447 | 4.087 | < 0.001 | -1.068 | 0.261 | -4.087 | < 0.001 |
| Latitude, Shp.PC1 | 0.033 | 1974.390 | 324.920 | 6.076 | < 0.001 | -49.410 | 8.130 | -6.077 | < 0.001 |
| Latitude, Shp.PC2 | 0.008 | -551.039 | 183.661 | -3.000 | 0.003 | 13.789 | 4.596 | 3.000 | 0.003 |
| Latitude, Shp.PC4 | 0.021 | -569.218 | 118.179 | -4.817 | < 0.001 | 14.243 | 2.957 | 4.817 | < 0.001 |
| Latitude, Shp.PC5 | 0.017 | 497.543 | 112.734 | 4.413 | < 0.001 | -12.450 | 2.821 | -4.414 | < 0.001 |
| Longitude, Prop.PC1 | 0.003 | 153.519 | 77.578 | 1.979 | 0.048 | 1.852 | 0.936 | 1.979 | 0.048 |
| Longitude, Prop.PC4 | 0.031 | 206.447 | 35.826 | 5.763 | < 0.001 | 2.491 | 0.432 | 5.763 | < 0.001 |
| Longitude, Prop.PC5 | 0.020 | -156.530 | 33.484 | -4.675 | < 0.001 | -1.888 | 0.404 | -4.675 | < 0.001 |
| Longitude, Shp.PC1 | 0.035 | 6981.260 | 1126.940 | 6.195 | < 0.001 | 84.220 | 13.600 | 6.195 | < 0.001 |
| Longitude, Shp.PC2 | 0.009 | -2041.937 | 637.037 | -3.205 | 0.001 | -24.634 | 7.685 | -3.205 | 0.001 |
| Longitude, Shp.PC4 | 0.024 | -2130.843 | 409.408 | -5.205 | < 0.001 | -25.707 | 4.939 | -5.205 | < 0.001 |
| Longitude, Shp.PC5 | 0.009 | 1275.300 | 392.930 | 3.246 | 0.001 | 15.390 | 4.740 | 3.246 | 0.001 |
| Year, Prop.PC4 | 0.011 | -93.845 | 26.449 | -3.548 | < 0.001 | 0.048 | 0.014 | 3.548 | < 0.001 |

We also find significant differences across latitude and longitude across many of the PCs. These differences may be driven by the large separation in recordings between Ottawa county and the other three counties –– indeed, nearly all of the same variables are identified (with the exception of Shp.PC3). Higher latitude and lower longitude recordings have lower recording lengths and lower bandwidth and power.

Lastly, Prop.PC4 is significantly correlated with year of recording. This suggests that bandwidth and power are what are predominantly changing as cowbird songs evolve. Whether this is due to actual changes in the songs or simply changes in recording technology cannot be determined with these data.

## Discussion

In this study, we have demonstrated that increased urban space has contributed to change in *M. ater* song over the last 50 or so years. Various habitats are associated with varying birdsong characteristics. For example, forested habitats are associated with low frequency and amplitude modulations as these characteristics favor reduced degradation by reverberation (Nicholls and Goldizen, 2006). In contrast, open habitats tend to favor songs that are of higher frequency and involve rapid modulation as there is limited reverberation and increased wind, which degrades birdsong (Henry and Lucas, 2010). Seeing as urban habitat differs significantly from natural habitat in terms of the degree to which auditory information is degraded by various factors, such as reverberation or distortion, it is expected that song from birds in urban environments differs from song of rural conspecifics. Background noise in cities, such as noise from traffic, tends to mask birdsong and interferes with the detection of acoustic signals by reducing the active space of birdsong (Nemethet al. 2013). Additionally, urban environments have higher reverberation than rural environments, further reducing the accurate transmission of birdsong (Warrenet al. 2006). In response to this background noise, birds tend to sing at higher frequencies to reduce attenuation of their song (Phillipset al. 2020). Seeing as there are observable differences in birdsong of urban and non-urban conspecifics, it is expected that there has been observable change in birdsong over time associated with the increased urbanization that has taken place over the last half-century.

In both humans and other species, there is evidence to suggest that behavior and skills can be transmitted between generations in what is thought to be synonymous with culture (Mace and Jordan, 2011; Whiten and Mesoudi, 2008). As with biological evolution, these cultural phenomena can change over time through selection on variation as an adaptation to changing environments in a process aptly named cultural evolution (Mundinger, 1980). The most robust evidence for the occurrence of cultural evolution outside of humans is in birds, in the form of birdsong (Whitenet al. 2017). Selective pressures that drive evolution of birdsong are likely those that promote vocalizations that best transmit through the bird’s habitat, in concordance with acoustic adaptation theory (Brown and Handford, 1996). Given this, two environmental factors appear to be the primary drivers of birdsong evolution: background noise and habitat structure (Phillips et al. 2020).

We found significant differences in both song properties and syllable shapes in *M. ater* song across Ohio counties. Song properties of recording length (Prop.PC1) and bandwidth and power (Prop.PC4) differed significantly across counties. All song shape principal components (Shp.PC1-5) differed significantly across counties. Most differences include either Ottawa or Franklin counties. This is sensible, as habitat structure is expected to differ greatly in these two counties. Ottawa County is geographically distant from the other three counties in which data was collected and sits against Lake Erie. Franklin County contains Ohio’s capital city, Columbus, and is therefore the most urbanized of the four counties. This is consistent with previous literature findings that suggest variation in habitat structure promotes different song characteristics, such as shifts in frequency or amplitude.

Further suggesting change in *M. ater* song in response to urbanization, is the correlation between song characteristics and cropland. It was found that change in cropland has a large impact on song properties and syllable shape, specifically Prop.PC1, Shp.PC1, Shp.PC2, Shp.PC4, and Shp.PC5. This indicates that song differs in regions that have gained or lost cropland from song in regions that have not had dramatic changes to amount of cropland. Seeing as a decrease in cropland in a geographic area is associated with an increase in urbanization in that same area, these data are congruent with our hypothesis that urbanization influences alterations in *M. ater* song. Additionally, the association between change in birdsong and change in cropland prevalence provides evidence for how direct anthropogenic habitat modification has impacted birdsong. Further, there were no significant correlations between *M. ater* song and other urbanization metrics.

Additional differences in birdsong were observed that cannot be confidently attributed to urbanization. The first of which is differences in songs seen across months of the year. We found that variation in duration and frequency (Prop.PC3) and bandwidth and power (Prop.PC4) of song was associated with month. Shp.PC3, Shp.PC4, and Shp.PC5 were also found to be associated with month. These associations could potentially be explained by differences in breeding effort throughout the year; *M. ater* breeding runs from April to August but peaks from May to June (Lowther 2020). Longitude and latitude are also associated with significant differences across principal components, with higher latitude and lower longitude recordings being associated with lower recording lengths and lower bandwidth and power. This is likely a result of Ottawa County being geographically distant from the other three counties. Finally, we observed a positive correlation between power and bandwidth (Prop.PC4) and year. This could indicate that these song properties are the predominant song characteristics that are changing with song evolution, however, it is also possible that this correlation is resulting from changes in recording technology and is not actually a result of true song property change over time.

Given the uncertainty regarding the cause of observed differences in cowbird song across months and years, further research is warranted to address these uncertainties. Additionally, at this time, the extent to which urbanization contributes to cultural evolution in *M. ater* compared to other drivers of cultural evolution remains unclear. Further investigative efforts regarding how urbanization interplays with other potential contributors to birdsong evolution, such as habitat fragmentation or population size, is warranted. Generalizability of these results to a larger group of organisms is also limited, given the restriction to one species. As such, examining our reported relationships in other species, such as non-brood parasites, may be a worthwhile avenue of research.

In examining the relationship between changes in *M. ater* song and habitat alteration over the last half-century, we were able to gather evidence favoring urbanization as a facilitator of brown-headed cowbird song evolution—specifically the observed differences in both song properties and shapes across Ohio counties as well as correlation between song characteristic change and change in cropland distribution. This observed association between change in *M. ater* song and increased urbanization provides increased insight into how anthropogenic habitat alteration influences cultural evolution in brood parasites. Seeing as birdsong plays a large role in ecological dynamics, such as mating and predation, variation in song resulting from anthropogenic change has the potential to drive speciation and other ecological change. The results of our analysis contribute to an increased understanding of how anthropogenic activity has altered avian behavior and suggest its continued influence on cultural evolution.

## Acknowledgements

We thank B. C. Carstens for helpful input. KLP was funded by NSF DEB-2016189.

## Appendix

**Appendix Table A1:** Non-significant Anova results comparing song metrics to spatiotemporal metrics. Response indicates the song metric evaluated, with “Prop.PCX” indicating a principal component calculated on the extracted song metrics and “Shp.PCX” indicating the same calculated on the SoundShape metrics. Predictor is the spatiotemporal measure compared. Other columns are degrees of freedom, sum of squares, mean squares, F value, and P value, respectively.

| Response | Predictor | DF | SSQ | MSQ | F | P |
| --- | --- | --- | --- | --- | --- | --- |
| Prop.PC2 | County | 3 | 38 | 12.53 | 0.969 | 0.40 |
| Prop.PC3 | County | 3 | 38 | 12.79 | 2.135 | 0.094 |
| Prop.PC5 | County | 3 | 24 | 7.983 | 2.348 | 0.071 |
| Prop.PC1 | Year | 1 | 30 | 29.68 | 1.65 | 0.19 |
| Prop.PC2 | Year | 1 | 35 | 34.99 | 2.709 | 0.10 |
| Prop.PC3 | Year | 1 | 6 | 5.626 | 0.936 | 0.33 |
| Prop.PC5 | Year | 1 | 0 | 0.435 | 0.127 | 0.72 |
| Shp.PC1 | Year | 1 | 10236 | 10236 | 2.61 | 0.10 |
| Shp.PC2 | Year | 1 | 20 | 19.5 | 0.016 | 0.90 |
| Shp.PC3 | Year | 1 | 143 | 143.0 | 0.215 | 0.64 |
| Shp.PC4 | Year | 1 | 10 | 9.7 | 0.019 | 0.89 |
| Shp.PC5 | Year | 1 | 27 | 26.6 | 0.057 | 0.81 |
| Prop.PC1 | Month | 1 | 31 | 30.59 | 1.701 | 0.19 |
| Prop.PC2 | Month | 1 | 8 | 7.634 | 0.59 | 0.44 |
| Prop.PC5 | Month | 1 | 0 | 0.393 | 0.115 | 0.73 |
| Shp.PC1 | Month | 1 | 387 | 387 | 0.098 | 0.75 |
| Shp.PC2 | Month | 1 | 4156 | 4156 | 3.407 | 0.065 |

**Appendix Table A2:**
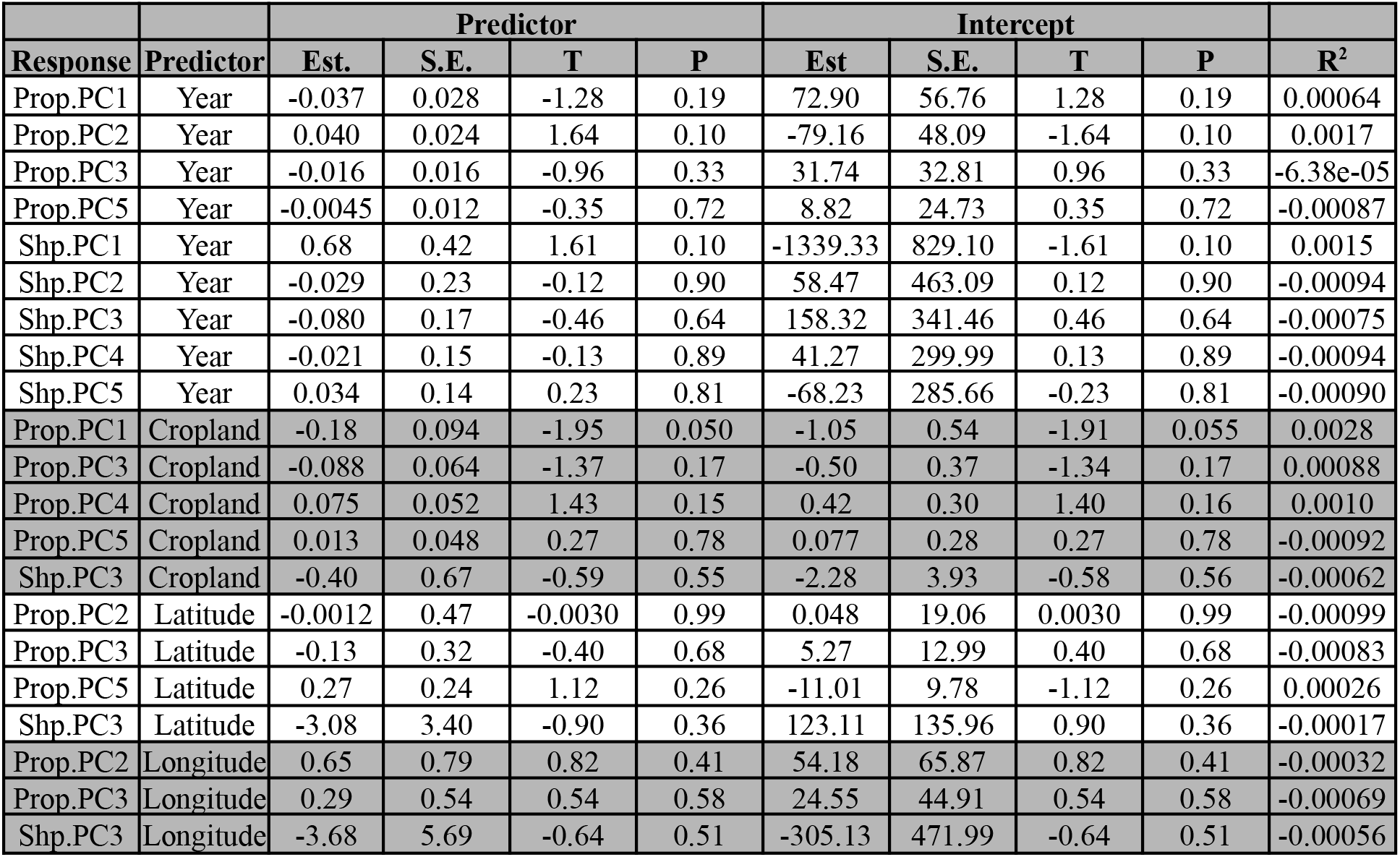
Non-significant Linear model results comparing song metrics to spatiotemporal metrics. Response indicates the song metric evaluated, with “Prop.PCX” indicating a principal component calculated on the extracted song metrics and “Shp.PCX” indicating the same calculated on the SoundShape metrics. Predictor is the spatiotemporal measure compared. Columns under the superheading “Predictor” give the linear model parameters for the predictor variable, while columns under the superheading “Intercept” give those parameters for the intercept variable. “Est” = estimated coefficient. “S.E” = standard error. “T” = T value. “P” = P value.

| Response | Predictor | Predictor |  |  |  | Intercept |  |  |  | R<sup>2</sup> |
| --- | --- | --- | --- | --- | --- | --- | --- | --- | --- | --- |
|  |  | Est. | S.E. | T | P | Est | S.E. | T | P |  |
| Prop.PC1 | Year | -0.037 | 0.028 | -1.28 | 0.19 | 72.90 | 56.76 | 1.28 | 0.19 | 0.00064 |
| Prop.PC2 | Year | 0.040 | 0.024 | 1.64 | 0.10 | -79.16 | 48.09 | -1.64 | 0.10 | 0.0017 |
| Prop.PC3 | Year | -0.016 | 0.016 | -0.96 | 0.33 | 31.74 | 32.81 | 0.96 | 0.33 | -6.38e-05 |
| Prop.PC5 | Year | -0.0045 | 0.012 | -0.35 | 0.72 | 8.82 | 24.73 | 0.35 | 0.72 | -0.00087 |
| Shp.PC1 | Year | 0.68 | 0.42 | 1.61 | 0.10 | -1339.33 | 829.10 | -1.61 | 0.10 | 0.0015 |
| Shp.PC2 | Year | -0.029 | 0.23 | -0.12 | 0.90 | 58.47 | 463.09 | 0.12 | 0.90 | -0.00094 |
| Shp.PC3 | Year | -0.080 | 0.17 | -0.46 | 0.64 | 158.32 | 341.46 | 0.46 | 0.64 | -0.00075 |
| Shp.PC4 | Year | -0.021 | 0.15 | -0.13 | 0.89 | 41.27 | 299.99 | 0.13 | 0.89 | -0.00094 |
| Shp.PC5 | Year | 0.034 | 0.14 | 0.23 | 0.81 | -68.23 | 285.66 | -0.23 | 0.81 | -0.00090 |
| Prop.PC1 | Cropland | -0.18 | 0.094 | -1.95 | 0.050 | -1.05 | 0.54 | -1.91 | 0.055 | 0.0028 |
| Prop.PC3 | Cropland | -0.088 | 0.064 | -1.37 | 0.17 | -0.50 | 0.37 | -1.34 | 0.17 | 0.00088 |
| Prop.PC4 | Cropland | 0.075 | 0.052 | 1.43 | 0.15 | 0.42 | 0.30 | 1.40 | 0.16 | 0.0010 |
| Prop.PC5 | Cropland | 0.013 | 0.048 | 0.27 | 0.78 | 0.077 | 0.28 | 0.27 | 0.78 | -0.00092 |
| Shp.PC3 | Cropland | -0.40 | 0.67 | -0.59 | 0.55 | -2.28 | 3.93 | -0.58 | 0.56 | -0.00062 |
| Prop.PC2 | Latitude | -0.0012 | 0.47 | -0.0030 | 0.99 | 0.048 | 19.06 | 0.0030 | 0.99 | -0.00099 |
| Prop.PC3 | Latitude | -0.13 | 0.32 | -0.40 | 0.68 | 5.27 | 12.99 | 0.40 | 0.68 | -0.00083 |
| Prop.PC5 | Latitude | 0.27 | 0.24 | 1.12 | 0.26 | -11.01 | 9.78 | -1.12 | 0.26 | 0.00026 |
| Shp.PC3 | Latitude | -3.08 | 3.40 | -0.90 | 0.36 | 123.11 | 135.96 | 0.90 | 0.36 | -0.00017 |
| Prop.PC2 | Longitude | 0.65 | 0.79 | 0.82 | 0.41 | 54.18 | 65.87 | 0.82 | 0.41 | -0.00032 |
| Prop.PC3 | Longitude | 0.29 | 0.54 | 0.54 | 0.58 | 24.55 | 44.91 | 0.54 | 0.58 | -0.00069 |
| Shp.PC3 | Longitude | -3.68 | 5.69 | -0.64 | 0.51 | -305.13 | 471.99 | -0.64 | 0.51 | -0.00056 |

